# TF-TWAS: Transcription-factor polymorphism associated with tissue-specific gene expression

**DOI:** 10.1101/405936

**Authors:** Yi-Ching Tang, Assaf Gottlieb

**Author notes:** Corresponding author: Assaf Gottlieb, Center for Precision Health, School of Biomedical Informatics, University of Texas Health Science Center in Houston, 7000 Fannin st, Houston, TX, 77030.

## Abstract

Transcriptional regulation is associated with a broad range of diseases. Methods associating genetic polymorphism with gene transcription levels offer key insights for understanding the transcriptional regulation plan. The majority of gene imputation methods focus on modeling polymorphism in the *cis* regions of the gene, partially owing to the large genetic search space. We hypothesize that polymorphism within transcription factors (TFs) may help explain transcription levels of their transcribed genes.

Here, we test this hypothesis by developing TF-TWAS: imputation models that integrate transcription factor information with transcription-wide association study methodology. By comparing TF-TWAS models to base models that use only gene *cis* information, we are able to estimate possible mechanisms of the TF polymorphism effect – TF expression or binding affinity within four tissues – whole blood, liver, brain hippocampus and coronary artery.

We identified 48 genes where the TF-TWAS models explain significantly better their expression than *cis* models alone in at least one of the four tissues. Sixteen of these genes are associated with various diseases, including cancer, neurological, psychiatric and rare genetic diseases. Our method is a new expansion to transcriptome-wide association studies and enables the identification of new associations between polymorphism in transcription factor and gene transcription levels.

## Introduction

Transcription regulation plays a critical role in cellular states and misregulation has been associated with a broad range of diseases (Lee and Young 2013). Several approaches have used genetic information in order to elucidate the association between transcription levels and phenotypic traits. Such approaches include association of phenotypes with expression quantitative trait loci (eQTLs) (Nica and Dermitzakis 2013) and more recently transcriptome-wide association studies (TWAS), including the PrediXcan method (Gamazon et al. 2015; Gusev et al. 2016), which develop models based on combinations of genetic variants for imputing gene expression prior to associating these imputed gene expression profiles with phenotypes. TWAS methods typically impute genes using only variants in *cis* with the gene whose expression they model, as the search space for possible *trans* associations is large and is thus harder to reach statistical significance.

Transcription factors (TFs) play important role in regulation of transcription by binding to *cis* regulatory regions in the vicinity of the regulated genes (Hobert 2008). Notably, polymorphism in TFs have recently been associated with several diseases including hypertension, coronary artery disease, Type 2 diabetes mellitus and lipedema (Fujimaki et al. 2015; Hamed et al. 2016; Palizban et al. 2017). Here, we hypothesize that polymorphism within transcription factors may be associated with the transcription levels of their transcribed genes. Thus, we test whether incorporating TF information into gene expression imputation models can improve the TWAS *cis* models. We introduce an extension to the TWAS *cis* models, which we call TF-enriched transcriptome wide association studies (TF-TWAS).

We tested three hypotheses concerning the way polymorphism within TFs may be associated with transcription levels of their transcribed genes: (1) *cis*-eQTLs of the TF may affect its transcription level and subsequently the transcription levels of the transcribed gene (Figure 1A); (2) TF polymorphism can affect binding of the TF to binding sites of their transcribed genes (Figure 1B); or (3) a combination of the two hypotheses (Figure 1C). We compared these three models to a base model, which includes only the *cis* variants of the gene (the TWAS *cis* model). We constructed TF-TWAS models in four selected tissues: whole blood, liver, brain (hippocampus) and coronary artery. We identified 48 hit genes for which the TF-TWAS models significantly improved upon the TWAS *cis* models. Twenty-one hit genes have at least one SNP within the TF with higher model weight than any of the *cis* SNPs and sixteen of these genes are associated with different cancer types, Alzheimer Disease, Paranoid Schizophrenia and rare genetic diseases such as William Syndrome, suggesting further investigation in future works. The TF-TWAS code is available at https://github.com/TangYiChing/TF-TWAS.

**Figure 1.**
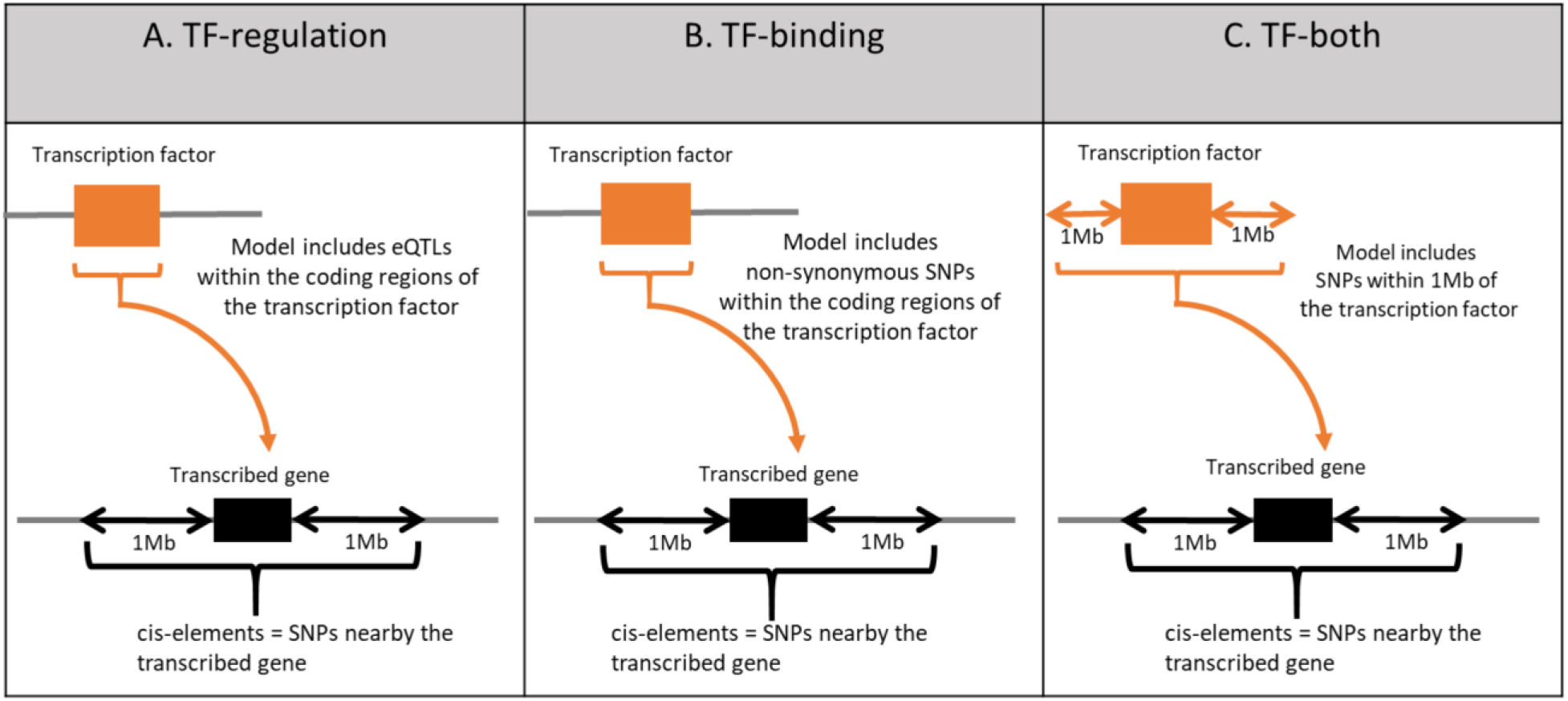
A illustration of the three tested hypotheses regarding the effect of TF SNPs on the expression of their transcribed genes. We test whether polymorphism in the TF (orange box) affect the transcription levels of the transcribed gene (black box) by testing the added effect of each TF-TWAS model relative to the baseline model. **TF-regulation model**: include eQTLs associated with the TF (A), **TF-binding model** includes non-synonymous SNPs within the associated TF boundary (B); and **TF-both model** include SNPs within 1MB of the associated TF coding region (C).

## Results

### Identifying TF-TWAS hit genes

Our objective was to identify genes whose gene expression imputation models were significantly improved by using information about polymorphism in their associated transcription factors, which we term TF-TWAS hit genes (see Methods). We computed the regression performance (R^2^) for a “base model”, based on the PrediXcan methodology {Gamazon, 2015 #125} (Methods), and the regression performance of three types of TF-TWAS models for 61,403 genes across four selected tissues: whole blood, liver, brain (hippocampus) and coronary artery. The three types of TF-TWAS models correspond to three proposed mechanisms of gene expression regulation: (1) effect of expression of the TF; (2) effect on the binding affinity of the TF to the TF binding site; and (3) effects that stem from both mechanisms. We will refer to these models as TF-regulation, TF-binding and TF-both, respectively (Methods).

Identification of TF-TWAS genes followed two steps: (1) comparing the regression performance of these three models to the base model; and (2) creating a background models for candidate outlier genes (see Methods, Table 1).

**Table 1.** Summary statistics of transcription factors and nearby SNPs.

|  | Whole blood | Liver | Brain hippocampus | Coronary artery |
| --- | --- | --- | --- | --- |
| # of samples | 177 | 60 | 54 | 72 |
| # of expressed genes | 22,423 | 21,167 | 22,836 | 23,354 |
| # of expressed gene with an associated TF | 14,635<br>(65.2%) | 15,056<br>(71.1%) | 15,953<br>(69.9%) | 15,759<br>(67.4%) |
| # TFs per gene | 6.3±8.3 | 6.7±8.4 | 6.5±8.2 | 6.3±8.2 |
| # of eQTLs per TF | 173.3±298.6 | 113.0±204.0 | 105.2±191.7 | 123.7±220.6 |
| # of nsSNP per TF | 1.3±2.3 | 1.3±2.3 | 1.3±2.3 | 1.3±2.3 |
| flanking TF SNPs | 6,732±1,468 | 6,735±1,471 | 6,733±1,473 | 6,732±1,473 |

\* nsSNP: non-synonymous SNP, eQTL: expression quantitative loci, flanking TF SNPs: SNPs within 1MB of TF.

In the base model comparison step, the regression scores (R^2^) of the TF-binding model obtained the highest correlation to the base model while the TF-both model obtained the least correlation in all four tissues (Figure 2). There were large variations in the correspondence of the three models to the base model in terms of regression performance across the tissues. While in whole blood all TF models where highly correlated to the base model (Pearson ρ>0.92), there were larger variations in other tissues, where liver showing the lowest correlation for all three models (Pearson 0.52<ρ<0.71) and brain showing the largest difference between TF models (Pearson 0.45<ρ<0.98).

**Figure 2.**
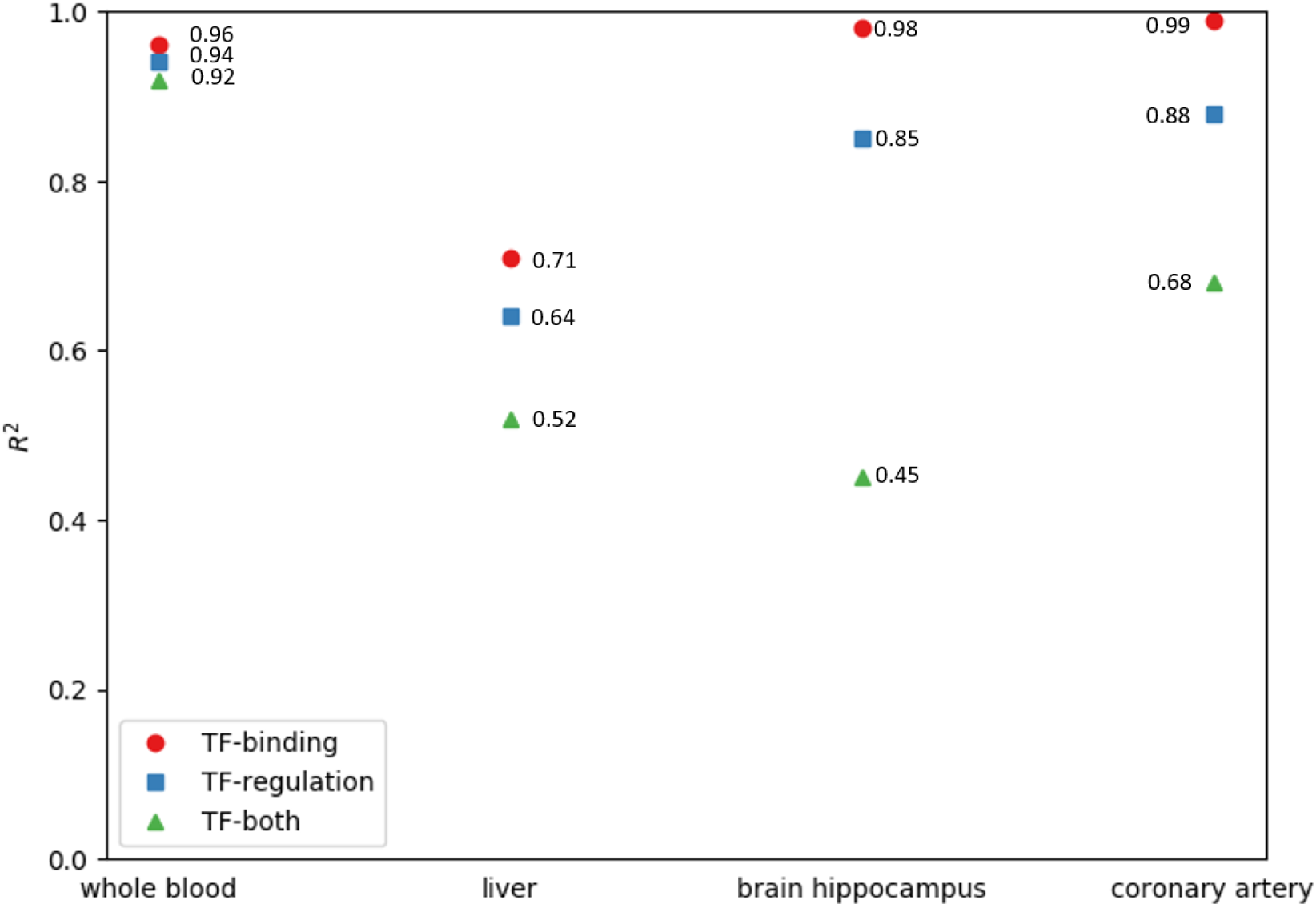
Comparison of prediction performance between the three TF-enriched models. Each marker represents the total variability between each the TF-enriched model and the baseline TWAS *cis* model across all genes and withint each of the four studied tissues.

Using the background model step, we identified one, thirty and seventeen significant genes for the TF-regulation, TF-binding and TF-both with false discovery rate of 0.01 within each tissue (Table S1). The majority of the genes from three models were detected in brain and coronary artery tissues (between one, four and six genes per tissue in all models and TF-binding is an exceptional model with twenty-five significant genes detected in brain), whereas we detected genes in liver tissue only for the TF-both model.

We found only single influential TF-SNPs per gene in the TF-binding model, while in the TF-both model the majority of the significant genes (nine out of twelve) genes have at least two influential TF-SNPs (see Methods, Table 2, Figure 3). As an extreme example, 28 influential TF-SNPs, distributed across nine TFs are associated with the HIST1H4J gene in the TF-both model. Some of the influential TFs were associated with more than two target genes. Two examples are GATA2, which is an influential TF in the expression of both ATG9B and AP3M1 genes, and YY1, which is an influential TF in the expression of both HIST1H4J and AP3M1 genes (Tables 2, S2-S3).

**Table 2.** List of genes with influential TF-SNPs. TFs that are associated with the same disease as their TF-hit genes are shown in bold.

| Gene | Influential TF | Associated diseases |
| --- | --- | --- |
| ATG9B | **ESR1, GATA2, NFKB1** | **Colorectal cancer (CRC)**; Essential Hypertension |
| ATXN1 | TRIP6 | Spinocerebellar ataxia type 1 (SCA1) |
| CPLX3 | TP53 | Chromosome 15Q24 Deletion Syndrome |
| GTF2I | ATF6, **CTCF**, EGR1, TS1, MIF, SP1, SRF, YBX1 | **Williams syndrome**; Chromosomal Deletion Syndrome. |
| HDC | KLF4, GATA2 | Tourette syndrome; Mast Cell Neoplasm |
| HSPA1B | **REST** | **Paranoid schizophrenia**; Acute mountain sickness |
| HSPE1 | HNF1A, DDIT3, PAX5, RXRA, POU3F1, RELA | Exocervical Carcinoma |
| IL15 | GATA2, HOXA10, **TP53** | **Rheumatoid Arthritis (RA)**; **Severe combined immunodeficiency (SCID)** |
| PJA2 | **ETS1** | **Non-small cell lung cancer (NSCLC)** |
| PSEN1 | **PAX5** | **Alzheimer disease**; Pick's disease of brain; Frontotemporal Dementia |
| PSMB3 | GABPA, GATA2, NFYA, **SP1**, YY1 | **Cystic Fibrosis (CF)** |
| RTN1 | CREB1, CUX1,RFX1 | Hereditary Spastic Paraplegia (HSP) |
| SETD2 | **E2F1** | **Gliomas**; Luscan-Lumish syndrome; Sotos Syndrome 1 |
| SHH | **CDC5L** | **Glioma**; Single median maxillary central incisor (SMMCI); Schizencephaly; Holoprosencephaly; Microphthalmia, isolated, with coloboma |
| SLC44A4 | GATA2, **RFX1** | **Deafness, Autosomal Dominant 72; Autosomal Dominant Non-Syndromic Sensorineural Deafness Type Dfna** |
| ZP1 | GATA2 | Oocyte maturation defect |
| AP3M1 | GATA2, YY1 | - |
| COQ10A | TFAP2C | - |
| HIST1H4J | ARNT, CTCF, HNF4A, MAX, MYC, SP1, STAT5, HNF4A, YY1 | - |
| PPRC1 | TAL1 | - |
| RING1 | XBP1 | - |

**Figure 3.**
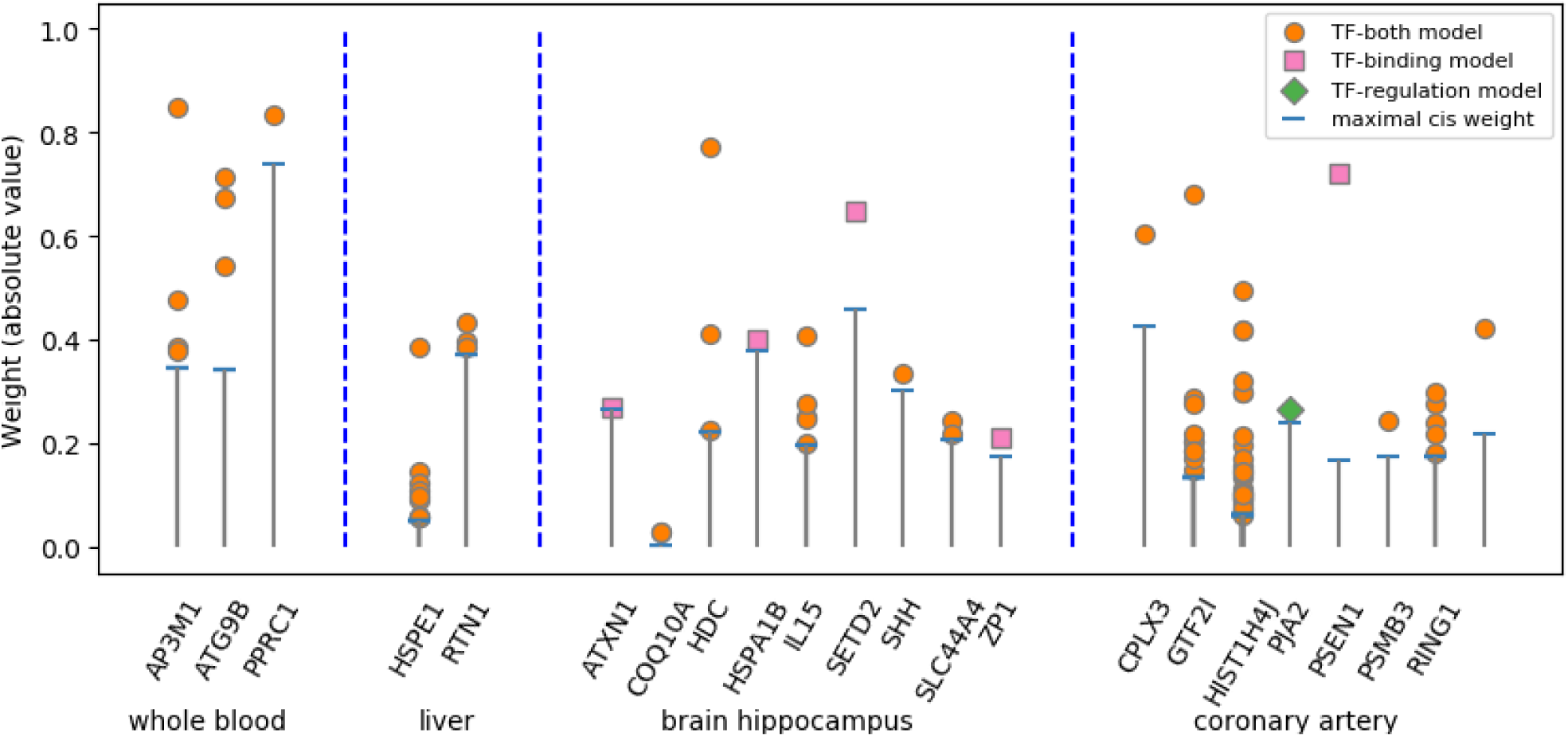
Strip plot of the list of influential TF-SNPs associated with TF-TWAS hit genes.

### TF-TWAS hit genes associated with disease

Our list of 48 TF-TWAS hit genes is enriched for two thyroid-related diseases, Hashimoto disease (seven hit genes in ToppGene Suite (Chen et al. 2009)) and Thyroid neoplasm (eight hit genes) (B&H FDR <7e^-4^). Three of the genes, BRAF, NFKB1 and AQP3, where found in both thyroid-related diseases. Interestingly, the set of TFs associated with these three genes is enriched for Thyroid Carcinoma (B&H FDR < 4e^-12^), out of which Fos proto-oncogene, AP-1 transcription factor subunit (FOS) is a TF associated with all three genes and high RNA levels of FOS were associated with Thyroid Carcinoma (Terrier et al. 1988).

We highlight two hit genes associated with pharmacogenomic traits. The first is APOA1,associated with response to fenofibrate in people with Hypertriglyceridemia (two genetic variants, rs964184 and rs2727786, (Aslibekyan et al. 2012)). Interestingly, one of its TFs, PPARA, has four other SNPs associated with decreased reduction in fasting IL-2 when treated with fenofibrate (Brisson et al. 2002; Frazier-Wood et al. 2013)). The second pharmacogenomic example is PPARD, associated with differential response to docetaxel and thalidomide in people with Prostatic Neoplasms (five different variants, (Deeken et al. 2010)). Two of its TFs, are also associated with these drugs, where RXRA show increased severity of Anemia when treated with docetaxel in people with Nasopharyngeal Neoplasmsa (Chew et al. 2014) and CTNNB1 is associated with increased response to thalidomide in people with Multiple Myeloma (Butrym et al. 2015).

We next focused on TF-TWAS hit genes with influential TFs - TF-TWAS hit genes that have at least one influential TF-SNP (Tables 2, S2-S3). We identified 21 disease-associated genes, associate with genetic diseases, mental and neuronal diseases (Tables 2, S2-S3). No genes were identified by more than one of the three TF models (Table S1). We highlight here two examples: For our first example, we identified a Presenilin 1 (PSEN1) as significant gene in coronary artery and furthermore, rs34810717 is an influential TF-SNP in its TF, Paired Box 5 (PAX5), with β weight of 0.7, while the maximal β of the *cis*-polymorphism of PSEN1 is 0.06 (Tables 2, S2-S3). Loss of function of PSEN1 gene has been correlated to early onset Alzheimer’s disease (Kelleher and Shen 2017). Both PSEN1 and its TF, PAX5, were reported to be associated with Alzheimer’s in African Americans and Caribbean Hispanics (Ghani et al. 2015).

Our second example involves gliomas, a type of tumor primary occurs in brain and spinal cord. We identified two TF-TWAS hit genes that are reported to be associated with gliomas along with and their corresponding influential TFs. They are the Sonic Hedgehog (SHH) gene and SET Domain Containing 2 (SETD2), identified in brain hippocampus according to the TF-both and TF-binding models, respectively (Tables 2, S2-S3). Aberrant SHH signal pathway and overexpression of its associated TF, CDC5L, are associated with tumor progression (Chen et al. 2016; Mariyath et al. 2018), while SETD2 and its influential TF, E2F1, are correlated with fast-growing gliomas (i.e., high-grade gliomas) (Yang et al. 2011; Fontebasso et al. 2013).

## Discussion

We hypothesized that polymorphism within transcription factors (TFs) may help model transcription levels of their transcribed genes. We tested our hypothesis using three types of TF-enriched models – a model focusing on TF expression, a model focusing on TF-gene binding and a model combining both. We found that different models identified different hit genes where the models better explained transcription levels than models using only *cis* SNPs, highlighting the complex regulatory mechanism underlying gene expression.

We observed large variations in the number of detected TF hit genes across TF models and across tissues. The largest number of TF-TWAS hit genes were identified by the TF-binding model, followed by the TF-both, while TF-expression model discovered only one hit gene. This result favors a mechanistic interpretation of the way polymorphisms in TF can affect gene expression by affecting their binding affinity over regulating the TF expression. We also observed large variations between the models in each tissue. In the hippocampus, we identified 25 hit genes using the TF-binding model and only six using the TF-Both, while in whole blood, the TF-Both model identified three genes and the TF-binding model only one.

As our results demonstrate, polymorphism in TFs affects only a relatively small fraction of genes (48 genes out of more than twenty thousand expressed genes in each tissue). This is not a surprising outcome, as TFs typically transcribe multiple genes. Changes in TF binding or expression that affect multiple genes may be detrimental to the individual and suffer from evolutionary pressure. Although our approach can only identify associations and not causal relations, they may still be useful to prioritize genes and TF pairs for further experimental validation.

We observed large variability in the number of discovered TF-TWAS hit genes across tissues and models. We could not establish a possible correlation between factors such as tissue sample size or the number of TF-associated SNPs used in the model with the number of TF-TWAS hit genes. We suggest that the lack of correlation with tissue sample size should be further examined using a larger set of tissues.

We have used curated sources for TF-gene associations, but they may still suffer from false associations and at the same time miss many true TF-gene associations. We assume that our two-stage approach to detect TF-TWAS hit genes is able to reduce the false hit genes to a minimum, but missing information about TF-gene associations probably reduced our ability to detect hit genes. Our hit genes list may thus be considered only partial to the true set of hit genes.

To summarize, we believe incorporating polymorphism in TFs is effective in modeling gene transcription levels for a subset of the human genes and detecting influential SNPs within TFs. Thus, these models should be considered in future gene expression imputation methods and provide a new approach for studying the genetic architecture of human disease.

## Methods

### Data

Genotype and expression data from the Genotype-Tissue Expression Project (GTEx) version 6 (Lonsdale et al. 2013) was retrieved from dbGaP. We imputed the GTEx genotype data using the Michigan imputation server (Das et al. 2016) using the Haplotype Reference Consortium r1.1 (McCarthy et al. 2016). Transcription factors and their transcribed genes were retrieved from Transcriptional Regulatory Relationships Unraveled by Sentence-based Text mining (TRRUST V2) (Han et al. 2017), the Human Transcriptional Regulation Interaction Database (HTRI) (Bovolenta et al. 2012) and the regulatory Network Repository of Transcription Factor and microRNA Mediated Gene Regulations (RegNetwork) (Liu et al. 2015). In total, we included 204,477 unique gene-TF pairs (with 6.3±8.3 TFs associated with each gene on average, Table 1). Genomic positions of the TFs were computed using the human genome assembly version 37 (GRCh37). For the TF-binding model, we used SnpEff v4.3 (Cingolani et al. 2012) for SNP functional annotation.

### Constructing TF-TWAS models

We compared the TF-TWAS models to a base model, which follows the models proposed by the TWAS algorithm PrediXcan (Zou and Hastie 2005). For the base model, we used the same parameters as PrediXcan, i.e. we used SNPs 1Mb from each of gene, filtered genotypes with imputation quality less than 0.8 and MAF < 0.01, adjusted for covariates including the first three principal components, 15 probabilistic estimation of expression residuals (PEER) factors (Stegle et al. 2012), gender and sequencing platform, and used elastic net regression.

For TF-TWAS models, we included the SNPs in the base model and TF-associated SNPs of all the TFs associated with transcription of the gene (Figure 1). We compared the base model to three sets of TF-associated SNPs, corresponding to different proposed mechanisms: (1) *cis*-eQTLs of the TF (association by regulation of the TF expression); (2) non-synonymous SNPs within the coding region of the TF (association by binding affinity); and (3) SNPs within 1Mb of the transcription factor coding region (both hypothesis). For brevity, we will refer to these models as TF-regulation, TF-binding and TF-both, respectively.

We compared these models across four tissues: whole blood (177 samples), liver (60 samples), hippocampus (54 samples) and coronary artery (72 samples) (Table 1). There were insignificant differences across tissues in the number of TFs associated with each gene, number of non-synonymous coding SNPs, eQTLs and SNPs within the 1Mb flanking regions of each TF (6.45, 0.9, 143 and 6733, respectively, Table 1).

### Identifying TF-TWAS hit genes

We compared the average regression coefficient (R^2^) over a 10-fold cross validation of the base model with the three types of TF models. We first detected candidate genes - genes whose TF model performance was higher than the base models by two standard deviations and Benjamini-Hochberg (Benjamini and Hochberg 1995) false discovery rate (B&H FDR) < 0.01. For each of these candidate genes, we constructed a background model to evaluate its significance.

Constructing the background models, we selected 100 random sets of TFs (each set with the same number of TFs as the true set of TFs associated with the gene) and computed the model R^2^ for each random set. Genes whose background model empirical p-values was less than or equal to a false discovery rate of 0.01 were identified as “TF-TWAS hit genes” (Figure 4).

**Figure 4.**
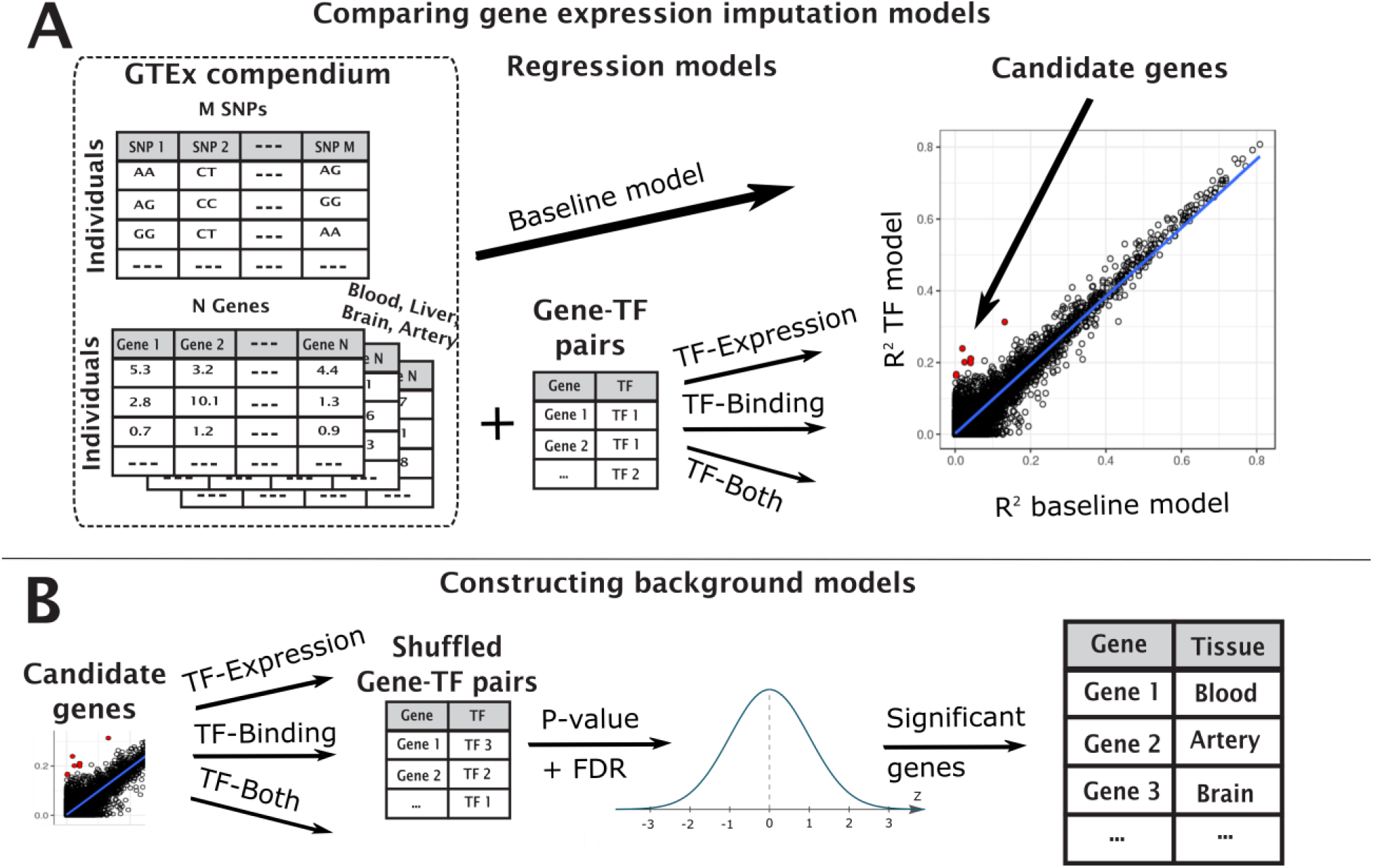
An illustration of the pipeline to identify TF-TWAS hit genes. We compute the baseline TWAS *cis* model and compare to one of the three TF-TWAS models to identify candidate genes (A). We compute a background model for each of the candidate genes to test their significance (B).

For the set of TF-TWAS hit genes, we further defined “influential TF-SNPs” as SNP within the TF region that obtain higher weights in the TF model than the maximal weight of its *cis* SNPs. We term their corresponding TFs “influential TFs”.

### TF-TWAS hit genes associated with disease

We identified associations between TF-TWAS hit genes with and diseases or pharmacogenomic traits using enrichment analysis through ToppGene Suite (Chen et al. 2009), searching online databases like OMIM (Hamosh et al. 2002), Malacards (Rappaport et al. 2013), PharmGKB (Whirl-Carrillo et al. 2012) and manual literature search.

## Data Access

GTEx data is available from dbGaP (accession phs000424.v7.p2). Transcription factors and their transcribed genes are publically available from Transcriptional Regulatory Relationships Unraveled by Sentence-based Text mining (TRRUST V2) (Han et al. 2017), the Human Transcriptional Regulation Interaction Database (HTRI) (Bovolenta et al. 2012) and the regulatory Network Repository of Transcription Factor and microRNA Mediated Gene Regulations (RegNetwork) (Liu et al. 2015).

## Acknowledgments

We would like to thank Arif Harmanci for constructive discussion and Jeffrey Chang for providing additional computing resources that allowed the heavy computations finish in a timely manner. The authors acknowledge the Texas Advanced Computing Center (TACC) at The University of Texas at Austin for providing HPC resources that have contributed to the research results reported within this paper.

## Author contributions

AG conceived the paper. YCT performed the experiments and analyzed the data. AG and YCT wrote the manuscript.

## Disclosure Declaration

The authors declare no conflict of interests.

## References

Aslibekyan S, Goodarzi MO, Frazier-Wood AC, Yan X, Irvin MR, Kim E, Tiwari HK, Guo X, Straka RJ, Taylor KD. 2012. Variants identified in a GWAS meta-analysis for blood lipids are associated with the lipid response to fenofibrate. PloS one 7:e48663.

Benjamini Y, Hochberg Y. 1995. Controlling the false discovery rate: a practical and powerful approach to multiple testing. Journal of the royal statistical society Series B (Methodological): 289–300.

Bovolenta LA, Acencio ML, Lemke N. 2012. HTRIdb: an open-access database for experimentally verified human transcriptional regulation interactions. BMC genomics 13:405.

Brisson D, Ledoux K, Bossé Y, St-Pierre J, Julien P, Perron P, Hudson TJ, Vohl M-C, Gaudet D. 2002. Effect of apolipoprotein E, peroxisome proliferator-activated receptor alpha and lipoprotein lipase gene mutations on the ability of fenofibrate to improve lipid profiles and reach clinical guideline targets among hypertriglyceridemic patients. Pharmacogenetics and Genomics 12:313–320.

Butrym A, Rybka J, Lacina P, Gebura K, Frontkiewicz D, Bogunia-Kubik K, Mazur G. 2015. Polymorphisms within beta-catenin encoding gene affect multiple myeloma development and treatment. Leukemia research 39:1462–1466.

Chen J, Bardes EE, Aronow BJ, Jegga AG. 2009. ToppGene Suite for gene list enrichment analysis and candidate gene prioritization. Nucleic acids research 37:W305–W311.

Chen W, Zhang L, Wang Y, Sun J, Wang D, Fan S, Ban N, Zhu J, Ji B, Wang Y. 2016. Expression of CDC5L is associated with tumor progression in gliomas. Tumour Biology: The Journal of the International Society for Oncodevelopmental Biology and Medicine 37:4093–4103.

Chew S-C, Lim J, Singh O, Chen X, Tan E-H, Lee E-J, Chowbay B. 2014. Pharmacogenetic effects of regulatory nuclear receptors (PXR, CAR, RXRα and HNF4α) on docetaxel disposition in Chinese nasopharyngeal cancer patients. European journal of clinical pharmacology 70:155–166.

Cingolani P, Platts A, Wang LL, Coon M, Nguyen T, Wang L, Land SJ, Lu X, Ruden DM. 2012. A program for annotating and predicting the effects of single nucleotide polymorphisms, SnpEff: SNPs in the genome of Drosophila melanogaster strain w1118; iso-2; iso-3. Fly 6:80–92.

Das S, Forer L, Schönherr S, Sidore C, Locke AE, Kwong A, Vrieze SI, Chew EY, Levy S, McGue M. 2016. Next-generation genotype imputation service and methods. Nature genetics 48:1284.

Deeken J, Cormier T, Price D, Sissung T, Steinberg S, Tran K, Liewehr D, Dahut W, Miao X, Figg W. 2010. A pharmacogenetic study of docetaxel and thalidomide in patients with castration-resistant prostate cancer using the DMET genotyping platform. The pharmacogenomics journal 10:191.

Fontebasso AM, Schwartzentruber J, Khuong-Quang D-A, Liu X-Y, Sturm D, Korshunov A, Jones DTW, Witt H, Kool M, Albrecht S et al. 2013. Mutations in SETD2 and genes affecting histone H3K36 methylation target hemispheric high-grade gliomas. Acta Neuropathologica 125:659–669.

Frazier-Wood A, Ordovas J, Straka R, Hixson J, Borecki I, Tiwari H, Arnett D. 2013. The PPAR alpha gene is associated with triglyceride, low-density cholesterol and inflammation marker response to fenofibrate intervention: the GOLDN study. The pharmacogenomics journal 13:312.

Fujimaki T, Oguri M, Horibe H, Kato K, Matsuoka R, Abe S, Tokoro F, Arai M, Noda T, Watanabe S. 2015. Association of a transcription factor 21 gene polymorphism with hypertension. Biomedical reports 3:118–122.

Gamazon ER, Wheeler HE, Shah KP, Mozaffari SV, Aquino-Michaels K, Carroll RJ, Eyler AE, Denny JC, Nicolae DL, Cox NJ. 2015. A gene-based association method for mapping traits using reference transcriptome data. Nature genetics 47:1091–1098.

Ghani M, Reitz C, Cheng R, Vardarajan BN, Jun G, Sato C, Naj A, Rajbhandary R, Wang L-S, Valladares O. 2015. Association of long runs of homozygosity with Alzheimer disease among African American individuals. JAMA neurology 72:1313–1323.

Gusev A, Ko A, Shi H, Bhatia G, Chung W, Penninx BW, Jansen R, De Geus EJ, Boomsma DI, Wright FA. 2016. Integrative approaches for large-scale transcriptome-wide association studies. Nature genetics 48:245.

Hamed WA, Hammouda GE, El-Hefnawy SM. 2016. Transcription factor 21 gene polymorphism in patients with coronary artery disease. Research Reports in Clinical Cardiology 55:13–18.

Hamosh A, Scott AF, Amberger J, Bocchini C, Valle D, McKusick VA. 2002. Online Mendelian Inheritance in Man (OMIM), a knowledgebase of human genes and genetic disorders. Nucleic acids research 30:52–55.

Han H, Cho J-W, Lee S, Yun A, Kim H, Bae D, Yang S, Kim CY, Lee M, Kim E. 2017. TRRUST v2: an expanded reference database of human and mouse transcriptional regulatory interactions. Nucleic acids research 46:D380–D386.

Hobert O. 2008. Gene regulation by transcription factors and microRNAs. Science 319:1785–1786.

Kelleher RJ, Shen J. 2017. Presenilin-1 mutations and Alzheimer’s disease. Proceedings of the National Academy of Sciences 114:629–631.

Lee TI, Young RA. 2013. Transcriptional regulation and its misregulation in disease. Cell 152:1237–1251.

Liu Z-P, Wu C, Miao H, Wu H. 2015. RegNetwork: an integrated database of transcriptional and post-transcriptional regulatory networks in human and mouse. Database 2015:bav095.

Lonsdale J, Thomas J, Salvatore M, Phillips R, Lo E, Shad S, Hasz R, Walters G, Garcia F, Young N. 2013. The genotype-tissue expression (GTEx) project. Nature genetics 45:580.

Mariyath MP, Shahi MH, Farheen S, Tayyab M, Khanam N, Ali A. 2018. Novel homeodomain transcription factor Nkx2.2 in the brain tumor development. Current Cancer Drug Targets doi:10.2174/1568009618666180102111539.

McCarthy S, Das S, Kretzschmar W, Delaneau O, Wood AR, Teumer A, Kang HM, Fuchsberger C, Danecek P, Sharp K. 2016. A reference panel of 64,976 haplotypes for genotype imputation. Nature genetics 48:1279.

Nica AC, Dermitzakis ET. 2013. Expression quantitative trait loci: present and future. Phil Trans R Soc B 368:20120362.

Palizban A, Rezaei M, Khanahmad H, Fazilati M. 2017. Transcription factor 7-like 2 polymorphism and context-specific risk of metabolic syndrome, type 2 diabetes, and dyslipidemia. Journal of research in medical sciences: the official journal of Isfahan University of Medical Sciences 22.

Rappaport N, Nativ N, Stelzer G, Twik M, Guan-Golan Y, Iny Stein T, Bahir I, Belinky F, Morrey CP, Safran M. 2013. MalaCards: an integrated compendium for diseases and their annotation. Database 2013.

Stegle O, Parts L, Piipari M, Winn J, Durbin R. 2012. Using probabilistic estimation of expression residuals (PEER) to obtain increased power and interpretability of gene expression analyses. Nature protocols 7:500.

Terrier P, Sheng Z, Schlumberger M, Tubiana M, Caillou B, Travagli J, Fragu P, Parmentier C, Riou G. 1988. Structure and expression of c-myc and c-fos proto-oncogenes in thyroid carcinomas. British journal of cancer 57:43.

Whirl-Carrillo M, McDonagh EM, Hebert J, Gong L, Sangkuhl K, Thorn C, Altman RB, Klein TE. 2012. Pharmacogenomics knowledge for personalized medicine. Clinical Pharmacology & Therapeutics 92:414–417.

Yang G, Zhang R, Chen X, Mu Y, Ai J, Shi C, Liu Y, Shi C, Sun L, Rainov NG et al. 2011. MiR-106a inhibits glioma cell growth by targeting E2F1 independent of p53 status. Journal of Molecular Medicine (Berlin, Germany) 89:1037–1050.

Zou H, Hastie T. 2005. Regularization and variable selection via the elastic net. Journal of the Royal Statistical Society: Series B (Statistical Methodology) 67:301–320.

